# The natural history of the black soldier fly, *Hermetia illucens*: insights from complete mitochondrial genome sequences

**DOI:** 10.1101/2021.10.21.465311

**Authors:** J Guilliet, G Baudouin, N Pollet, J Filée

**Affiliations:** Laboratoire Evolution, Génomes, Comportement, Ecologie CNRS Université Paris Sud UMR 9191, IRD UMR 247 CNRS Avenue de la Terrasse, Bâtiment 13, Boite Postale 1, 91198 Gif sur Yvette France

**Keywords:** *Hermetia illucens*, phylogeny, mitochondrial genome, phylogeography

## Abstract

The Black Soldier Fly (BSF) *Hermetia illucens* is a cosmopolitan fly heavily used by industrial companies to reduce biowaste and produce protein and fat for poultry and aquaculture feed. However, the natural history and the genetic diversity of the BSF is poorly known. In this study, we analysed 677 CO1 sequences derived from samples found all over the five continents leading us to the discovery of 52 haplotypes including 10 major haplotypes. We refined the definition of these haplotypes by sequencing 59 mitochondrial genomes. We could derive an estimate of the separation events of the different haplotypes at more than two million years for the oldest branches. This worldwide cryptic genetic and genomic diversity is mirrored at local scale in France in which we found five major haplotypes sometimes in sympatry. Our data resolve the phylogenetic relationships between the major lineages and give insights into the dispersal and the numbers of BSF neo-introduction at global and local scales. Our results indicate that the genetic and genomic diversity of commercial BSF stock is very low and these brood stock participated in the dissemination of the BSF in the wild. Taken together these results call for a better understanding of the genomic diversity of the BSF to unravel possible specific adaptations of the different lineages for industrial needs and to initiate the selection process.

## Introduction

The black soldier fly (BSF) *Hermetia illucens* (Linnaeus, 1758) is a Diptera of the Stratiomyidae family. It probably originated from Mexico, then subsequently spread to the south of the USA and across Latin America over the last thousand years (Hauser et al. 2015). These geographical areas are now considered as its natural area (Rozkosný, 1983,Hauser et al. 2015). The BSF is now a cosmopolitan fly that can be found in tropical, subtropical, and tempered regions, from the 40th parallel North and the 45th South (Rozkošný 1983 ; Mason et al, 2009) where it is considered as an exotic, non-invasive species (https://tinyurl.com/5353hvjv, Wang and Shelomi, 2017). This fly likely spread through shipping routes (Picker et al, 2002). While studies based on naturalistic observations show that it was first discovered in Malta in 1926 (Lindner, 1936), it has been suggested that it was introduced in Italy about 500 years ago (Benelli et al, 2014). The BSF appeared in the early 1950s in the South-East of France Barbier 1952 before spreading through the South-West, (Dauphin 2003) along the Rhône valley (Richoux 2009) and finally along the Atlantic coast (Maquart et al, 2020). Currently, the BSF can be found on the whole European territory with few exceptions. It is reasonable to assume that the rise of international traffic and the industrial use of the BSF may have led to a steady flow of introductions in France and all over the world (Leclerq 1969, Sheppard et al, 1994).

The BSF was initially used in forensic entomology to date the post-mortem time (Lord et al, 1994), now it is heavily used by industrial companies to reduce biowastes (Gold et al, 2020 ; Hoc et al, 2020), to produce protein (Hoc et al, 2020 ; Ewald et al, 2020 ;Giannetto et al, 2019) and fat (Hoc et al, 2020 ; Ewald et al, 2020 ; Spranghers et al, 2017). These products are respectively used to make animal food (Huis 2013) or biodiesel (Leong et al, 2016 ; Surendra et al, 2020).

BSF industries face similar issues as other farms: knowing how to master the biological and genetic aspects of their model to get the best out of it. To this end, it is important to know the genetic diversity of BSF existing internationally, nationally, and locally. A lot of articles exist about the nutritional aspect of the BSF (Spranghers et al, 2017 ; Heuel et al, 2021 ; Schiavone et al, 2019 ; Neumann et al, 2018 ; Biasato et al, 2019) or its microbiota (Khamis et al, 2020 ; Erickson et al, 2004 ; Liu et al, 2008), few have focused on the genetic diversity of the BSF. We have some evidence on the extent of BSF genetic diversity from previous studies. 10 haplotypes based on the CO1 gene were found in South Korea with 245 individuals (Park et al, 2017). Still on the CO1 gene, and on a worldwide sampling, 56 haplotypes were found with a divergence rate up to 4.9% and it has been shown that individuals belonging to divergent lineages are still interfertile (Ståhls et al 2020). Finally, based on 15 microsatellite markers, 16 genetic clusters were found with hot spots in South America (Kaya et al, 2021). Other works focused on more fundamental aspects of BSF and led to the assembly of a reference mitochondrial genome as well as a chromosomal-scale nuclear genome (Qi et al, 2017 ; Generalovic et al, 2020 ; Zhan et al, 2020).

Within the framework of the BSF industry, phylogeography can allow on the one side to account for genetic diversity at different scales (global, national, and local) and on the other side, to have keys on the geographical origin of the species present in a territory or used by companies. This study may suggest local adaptations or some phenotypic plasticity, due to habituation to different biomes.

In this study, we quantified this diversity at different scales: worldwide by studying the variation within the CO1 gene, and more locally at the level of France. We also analysed the role of livestock BSF farming in the diversity present on the French territory. Our report is based on data obtained by sequencing 113 individuals for the CO1 gene and also the mining of 564 CO1 sequences. We alsos sequenced and assembled 59 mitochondrial genomes. We also show the great and hidden genetic diversity within the species *Hermetia illucens* at all observation scales with a specific focus on France (90 French individuals were analysed, allowing us to find 9 different haplotypes). In addition, we are also providing here a date of the separation events of the different haplotypes found. This separation turns out to be more than two million years old for the most distant haplotypes.

## Materials and methods

### Sampling

We decided to carry out a global sampling followed by a specific focus on France. A detailed table of samples and associated metadata is provided in (SUPP_TABLE 001 (SUPP_MATERIAL)). The origin of all CO1 sequences used in our study is summarized in Table 1 and a summary of sample numbers by continent is provided in Table 2. Samples were collected during summer 2020 and stored in 100% ethanol at −20°C prior to DNA extraction. In France, the collection was done with the help of the compost citizen network (https://reseaucompost.org/). The larvae came exclusively from compost and the adults collected were close to the composting area. Between one and 10 individuals (larvae or adults) were collected per area. In some French localities, collections in the wild have been made near BSF breeding areas (about five kilometres). For the other parts of the world, the individuals came either from compost or from breeding farms.

**Table 1:** Origin of CO1 sequences and mitochondrial genomes

| ORIGIN | NUMBER OF INDIVIDUALS (CO1) | NUMBER OF INDIVIDUALS (MITOCHONDRIA) |
| --- | --- | --- |
| **COLLECTION FROM THE WILD** | 91 | 43 |
| **INDUSTRIAL ORIGIN** | 23 | 13 |
| **MINED FROM DATABASES (GENBANK<sup>39</sup> & BOLD<sup>40</sup>)** | 560 | 1 |
| **ASSEMBLED FROM SRA<sup>41</sup> (RAW SEQUENCES)** | 3 | 3 |
| **TOTAL** | 677 | 60 |

**Table 2:**
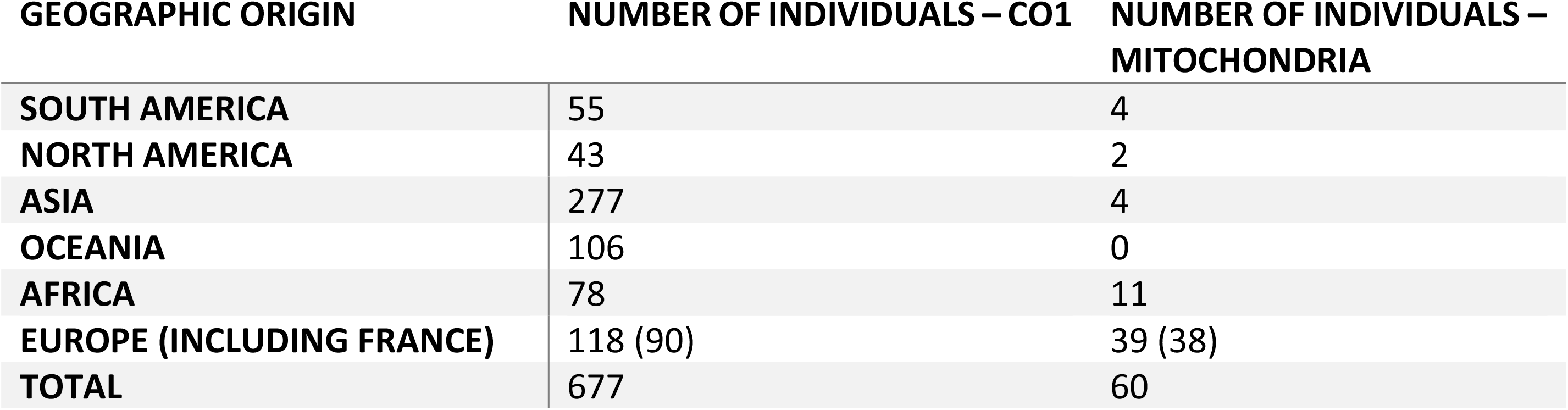
Geographic origin of CO1 sequences and mitochondrial genomes

| GEOGRAPHIC ORIGIN | NUMBER OF INDIVIDUALS – CO1 | NUMBER OF INDIVIDUALS – MITOCHONDRIA |
| --- | --- | --- |
| **SOUTH AMERICA** | 55 | 4 |
| **NORTH AMERICA** | 43 | 2 |
| **ASIA** | 277 | 4 |
| **OCEANIA** | 106 | 0 |
| **AFRICA** | 78 | 11 |
| **EUROPE (INCLUDING FRANCE)** | 118 (90) | 39 (38) |
| **TOTAL** | 677 | 60 |

We gathered all the CO1 sequences available on NCBI and Bold, as well as whole genome sequencing data in SRA, and reconstructed the mitochondrial genome. In France, sampling was carried out during the summer of 2020 in shared and personal compost sets. We obtained larvae in the areas of Blois, Bordeaux, Toulouse, Montpellier, Lyon, Paris. In Africa, individuals from compost or livestock farms were collected during 2019. In Latin America, larvae from French Guiana compost were collected during the summer of 2019. An adult found in 2018 in Mexico. In Asia, individuals collected from Taiwan either from a breeding farm or compost. These were mainly L5 larvae and pupae; some adults were also recovered. We had the surprise in the Bayonne area to have BSF-like larvae, which turned out to be *Exaireta spinigera* larvae.

### DNA extraction

We extracted DNA from larvae, pupae and imago under the same conditions by grinding individuals in liquid nitrogen using a mortar and pestle. The grinded samples were then processed according to the protocol of the Macherey-Nagel AXG 100 (https://tinyurl.com/46a7xejr) kit for the isolation of genomic DNA from tissue with some modification : we added a second purification step with 3.5 ml of freshly prepared ethanol and centrifugation at 15,000 rpm at 4 ° C for 15 min just before the final re-suspension. The DNA pellet was finally re-suspended in 100 μL of Tris 10 mM EDTA 1 mM pH 8.0. The purity was evaluated by spectrophotometry using a Nanodrop (Desjardins and Conklin 2010), and the concentration was quantified using the Qubit kit DNA Broad range (https://tinyurl.com/evaah44c). DNA integrity was monitored using routine agarose gel electrophoresis.

### CO1 Sequencing

We designed specific CO1 primers from the reference mitochondrial DNA (Qi et al, 2017) (SUPP_TABLE 002 (SUPP_MATERIAL)), then a mix was realized using NEB 5X standard buffer One Taq. The mix is then amplified in ONETAQ HOT 45 PCR. We purified the PCR products using the Illustra Exostar 1-step GE Healthcare kit before Sanger cycle sequencing. Cycle sequencing reactions were performed using the BigDye 3.1 chemistry, purified by precipitation and sequenced using an ABI 3130 sequencer (Applied Biosystems). Base calling was performed using the Sequencing analysis software version 5. 3 (Applied Biosystems). We assembled forward and reverse reads using the Geneious (https://www.geneious.com) assembler with the sensitivity set at medium/fast. The regions containing the primers were excised and we kept only the sequences having more than 75% of high quality basecalls.

### Multiple sequence alignment and haplotype network reconstruction

We aligned CO1 sequences obtained in the laboratory as well as those recovered from the databases using the MAFFT (Katoh and Standley 2013) aligner included as a plug-in in the Geneious (https://www.geneious.com) software. We used the algorithm G-INS-i with a score matrix 200 PAM / k=2, a gap open penalty at 1.53 and offset value at 0.123. We exported the multiple sequence alignment in nexus format (Maddison et al, 1997) for haplotype network reconstruction and visualization using POPART (Leigh and Bryant 2015). We modified the output file to add geographical origins to the sequences.

### Shotgun sequencing and mitochondrial DNA assembly

A total of 56 BSF individuals were used for whole-genome shotgun sequencing using an Illumina method by Novogene UK (Cock et al, 2010 ; Hansen et al, 2010 ; Erlich et al, 2008 ; Jiang et al, 2011 ; Yan et al, 2013). The technique used is pair-end with a coverage rate of 4,68X (SD 0,50). The resulting sequences were checked for quality and counted for read parity. We take advantage of the fact that mtDNA is highly repeated in insect DNA. In insects, mtDNA represents on average 0.42% of the total DNA in the genome sequence project (Meng et al, 2019). As stipulated, we used 5000 kb of data/sample (4-5× coverage) for the mitochondrial assembly, which in average contains (5000*0,42)/100 = 21000kb of mtDNA. As the mitochondrial genome is about 15 kb, we obtain a coverage of 1400× for this genome which is largely sufficient to obtain a complete genome in one contig. The sequences were assembled de novo with the MEGAhit (Dinghua et al, 2015) software (based options); the mitochondrial genomes were thus recovered from the assemblies by blast (Altschul et al, 1990) using the reference BSF mitochondrial genome sequence (NC_035232). Using MITOS online software (Bernt et al, 2013) the mitochondrial DNAs were annotated, and the D-Loop zone removed. To these sequences were added 4 complete mitochondrial sequences from databases or reconstructed via read sequences available on the SRA (Leinonen et al, 2011) using the same assembly procedure and then aligned under the same conditions as previously mentioned.

### CO1 and whole mitochondrial genome phylogeny

We use the MEGAX (Sudhir et al, 2018) software to determine the best model and then compute a ML Tree with 500 Bootstraps using a complete deletion parametre. We visualized the data with the online tool ITOL (Letunic and Bork, 2021), and we rooted the tree with the Stratiomyidae *Exaireta spinigera*. We chose *Exaireta spinigera* because it was the closest species for which we have a complete mitochondrial genome (the blastn result of the complete mitochondrial genome of *Hermetia illucens* vs *Exaireta spinigera* gives us 81.28% of similarity). To simplify the interpretation of the phylogeny, we have kept only a single identical sequence per haplotype.

### Time tree

We used the BEAST software (Suchard et al, 2018) to estimate the divergence time of the different BSF groups. The analysis was performed with an expansion growth model and an uncorrelated lognormal relaxed clock with the proposed insect molecular clock. 10 million iterations were performed and then with the TreeAnotator (Helfrich et al, 2018) the obtained trees were clustered with a burn of 25% of the trees. The visualization of the final tree was done with the online software ITOL13.

## Results

### Haplotype Network

We reconstructed an haplotype network from the analysis of 677 sequences spanning 658 nucleotides of the CO1 mitochondrial gene. A total of 52 distinct haplotypes were delineated from the analysis of this CO1 fragment that were made of one to 230 fly samples (Figure 1).

**FIG 1.**
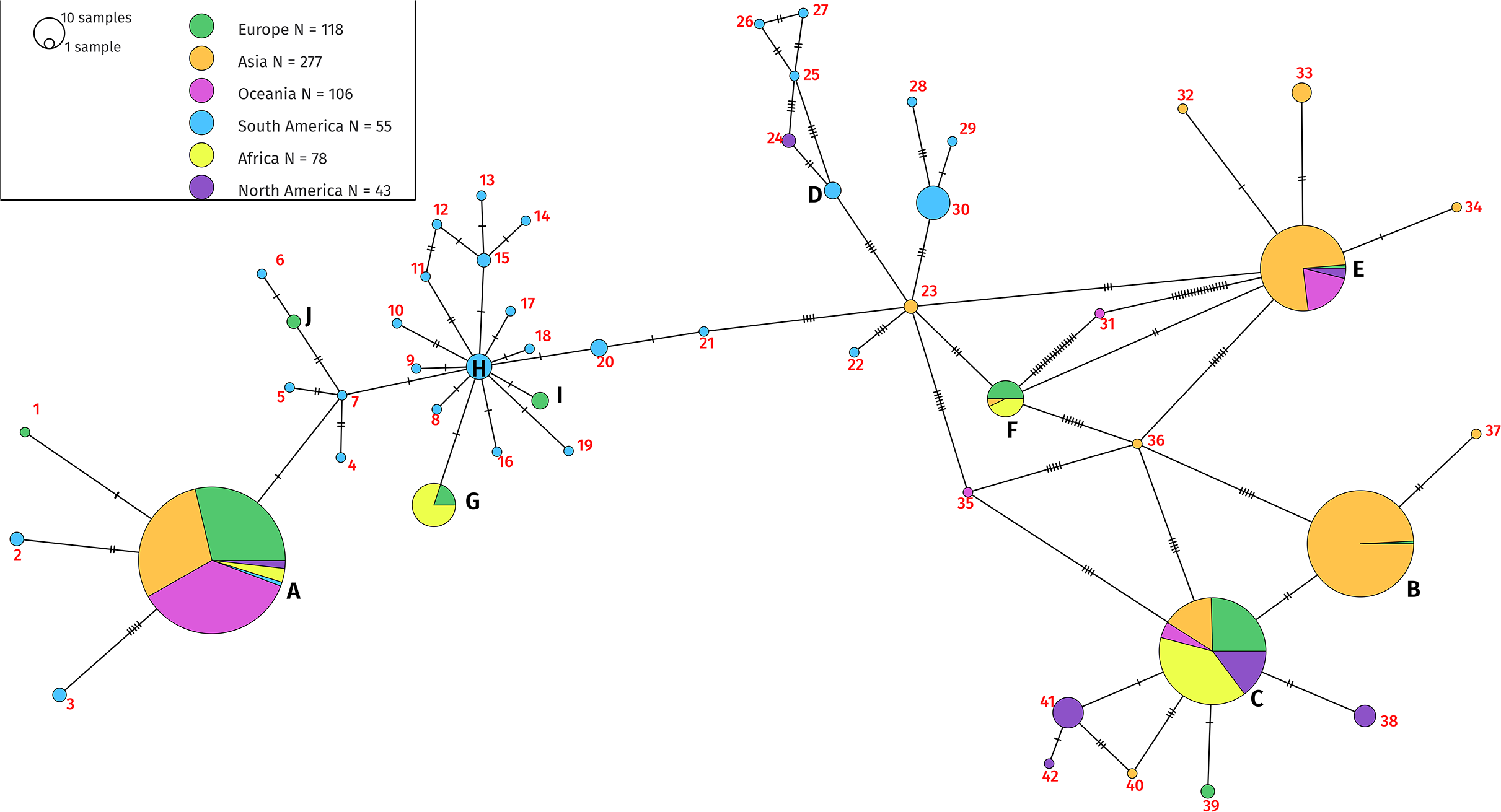
Haplotype network generated by PopArt after multiple alignment, based on the CO1 gene of Hermetia illucens. Analysis of the differences between the CO1 gene sequences of 677 individuals. The pie charts represent the same haplotype, and their relative size gives an indication of the number of individuals sharing this haplotype. The number of bars between each segment indicates the number of differences between the two sequences. Within a pie chart the colors represent the geographical origin of the individuals. The major haplotypes are those with more than five sequences (A B C E F G H 30 38 & 41). The minor haplotypes are the others. The list of sequences and their exact origin is available as an additional document.

We found 73 segregating sites with a Tajima-D of 45.1413 suggesting a lack of rare alleles, which could come from a contraction of the population.

There were 31 haplotypes made of a single sequence. We observed 11 haplotypes made of between two and four sequences derived from the same continent; that total of 42 haplotypes will be referred to as minor haplotypes. We observed 10 haplotypes made of between five and 230 sequences including 6 multi continental haplotypes that we will refer to as major haplotypes. In conclusion, we found an important diversity both within and between continents using the CO1 phylogenetic mitochondrial gene. Minor haplotypes represent 8.5% of all sequences (58 sequences) and 80.8% of haplotypes. Major haplotypes represent 91.5% of the sequences (619 sequences) and 19.2% of the haplotypes.

In Europe, we found 10 distinct haplotypes from 118 sequences, nine for Asia from 277 sequences, five for Oceania from 106 sequences, 30 for South America from 55 sequences, four in Africa from 78 sequences and seven in North America from 43 sequences. Some major haplotypes are represented in a more restricted area, for example haplotype B is mainly present in Asia (figure 1). We observed a high haplotype diversity in South America with 28 minor haplotypes that contain between one and four sequences. The haplotype C contains all the sequences derived from farmed strains of the industry working on BSF.

### Global distribution analysis

We visualized the actual distribution of the different haplotypes found previously on a distribution map (figure 2).

**Fig 2.**
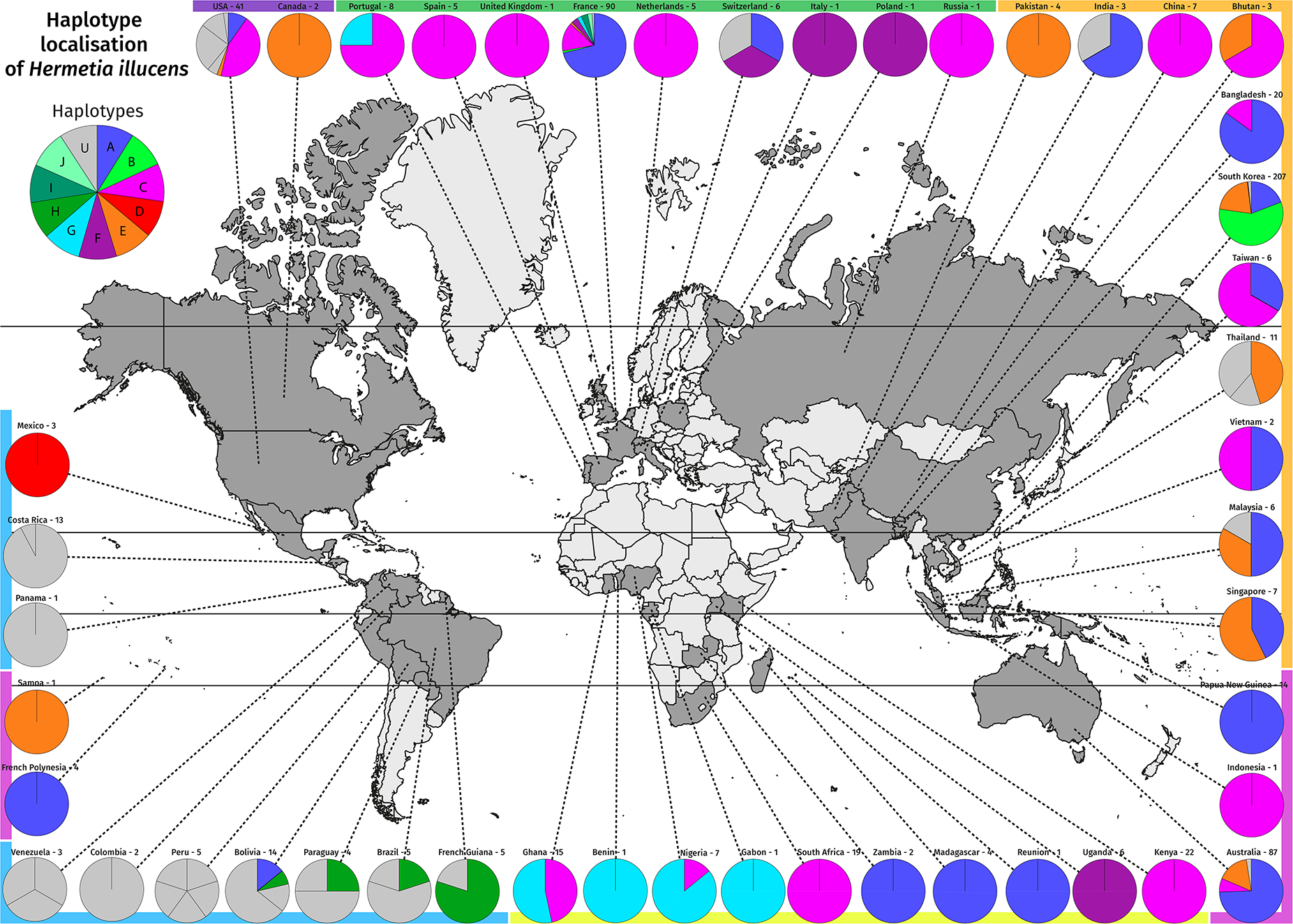
Global distribution of the different *Hermetia illucens* haplotypes based on the CO1 gene. The pie charts represent the proportion of each haplotype within a country. The name of the country is followed by the number of individuals analysed in that country. A colored band representing the continents is next to each pie chart. The colors represent the haplotypes determined in figure 1. The grey haplotypes are those that were represented with a number in figure 1. A separation is made in the pie chart when the haplotypes change for the same country. For example, Venezuela contains three unique haplotypes, meaning that these three individuals are different from each other. A summary table can be found in the supplementary table 3.

Each of the haplotypes from A to J is now represented with a color; haplotypes that are numbered in the haplotype network (figure 1) are now represented in grey.

We observed a great variability in South America highlighted on the map by the grey pie charts representing mostly minor haplotypes. We observe 22 singletons in South America out of the 52 haplotypes found in total, which is more than 40% of the total diversity. On the other hand, we found six out of ten major haplotypes in Europe.

Haplotype A and C were the most represented haplotypes in our sampling: Haplotype A was present on all continents, and represented 34% of all sequences, and haplotype C was only absent from South America, and represented 18% of all sequences. Some haplotypes seemed to be restricted to certain regions of the world, such as haplotype G which was present in all the localities in West Africa and surprisingly in a Portuguese sample, or the haplotype H in South America. Haplotype F was present in Europe, Uganda, and South Korea. Haplotype E was more present in Asia and Oceania as well as in North America. Finally, some haplotypes such as B, I and J present rare sequences found in more restricted areas such as B in Korea or H and J in France (figure 2).

### CO1 Phylogeny

During our CO1 gene analyses, some BSF-like larvae were identified as *Exaireta spinigera* larvae. The sequences were amplified and sequenced with the same primers as for BSF.

By taking only one sequence for each haplotype we have reconstructed a phylogeny of the CO1 gene sequence. The CO1 phylogeny was rooted with the *Exaireta spinigera* (figure 3). A sample from Cameroon was revealed to belong to a closely unidentified species of BSF and was termed *Hermetia sp*. Cameroon in our subsequent analysis (blastn score between *Hermetia* sp. from Cameroon and BSF gives us 85.01% identity with an E Value of 2e-174).

**FIG 3.**
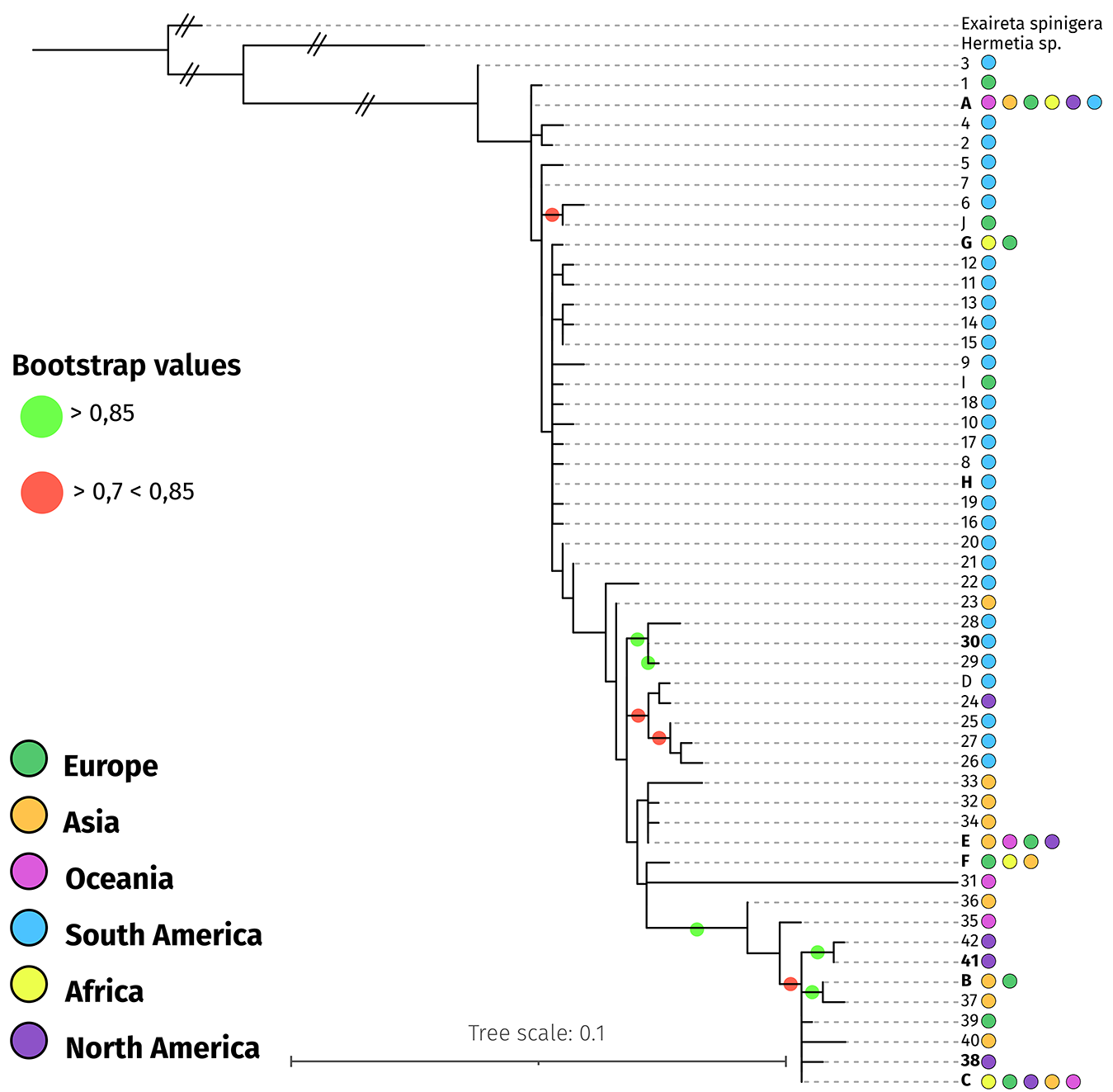
ML phylogenetic tree. Rooted phylogeny with *Exaireta spinigera* (Stratiomyidae). Bootstrap values are materialized with colored circles: green for values > 0.85 and red for values between 0.7 and 0.85. We only kept one sequence per haplotype to visualise the relationship between them. Next to each haplotype, colored circles represent the geographical origin of the species present in this haplotype. The order indicates the relative abundance of the individuals of a container in each haplotype. The statistical support was estimated with 500 Bootstraps with MegaX software using the Tamura 92 G+I model.

The CO1 gene phylogeny supports the analyses performed with the haplotype network (Figure 1), showing remarkable sequence diversity among the CO1 gene. Interestingly, all the commercial strains are found in the C haplotype (a detailed phylogenetic tree is given as a supplementary with an asterisk placed on individuals from farms - (SUPP_Figure 003 (SUPP_MATERIAL))).

A group containing haplotypes 35 to 42 and haplotypes B and C forms a divergent group from the others, with strong statistical support (bootstrap value of 0.99). Some sequences of individuals captured in the wild have been found near BSF commercial breeding sites, for example, we have captured C haplotype individuals in Avignon close by companies that are working with the BSF.

Although the global resolution of the deep branches of the tree is low, it’s also possible to better understand the relationship between the haplotypes: haplotype B and C are close to each other and well separated from other haplotypes. The other haplotypes are not very well resolved with the CO1 sequences. Finally, the samples coming from Latin America are scattered all over the tree, showing their great diversity and supporting the hypothesis of the geographical origin of the BSF.

### The global BSF diversity is mirrored at the local level in France

Among the 10 major haplotypes, our sampling effort in France showed that five are present in France in addition to a unique haplotype only found in France (Figure 4) in La Rochelle. Haplotype A remains the dominant haplotype in France (71.1% of all sequences) in seven of the eight localities (excluding Beaufort en Vallée where the flies come from a company). Regarding the haplotype C, BSF belonging to this group have been found in four locations in the regions of Poitiers, Bordeaux, Beaufort-En-Vallée and Montpellier. Except for Beaufort-En-Vallée, where the flies come from a company, the other BSF come from natural environments. BSF industries exist in all these localities. More surprisingly, different haplotypes coexist in some identical localities. For example we found three different haplotypes in the region of Paris and four in the region of Montpellier. Finally, we found a singular haplotype in Colomiers (J) and La Rochelle (1) .

**FIG 4.**
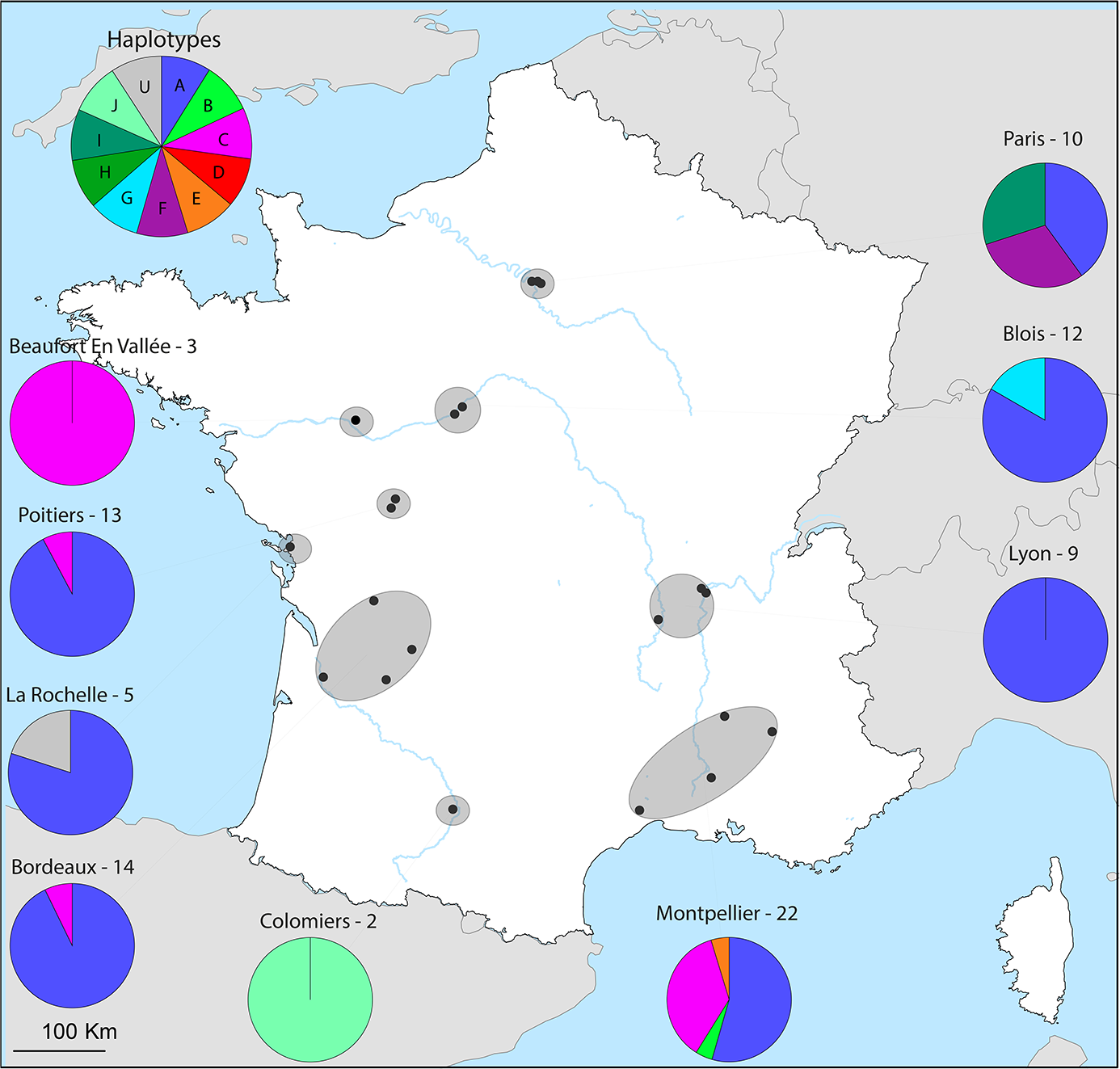
Distribution of the different haplotypes of *Hermetia illucens* in France. The dots represent the precise area of collect. The pie charts represent the distribution of haplotypes in the corresponding area, the names of the main cities in each area are indicated above as well as the number of sequenced individuals. Paris includes Vincennes and Nogent Sur Marnes. Blois includes Valloire Sur Cisse. Lyon includes Saint Marcellin En Forez, Corbas, Champagne Au Mont d’Or and Dardilly. Montpellier includes Avignon, Trescléoux and Dieulefit. Bordeaux includes Lormont La Force and Périgueux. Poitiers includes Saint Georges Lès Baillargeaux.

### Complete mitochondrial genome analyses

To better understand BSF phylogeny and to estimate the timing of haplotype divergence, we reconstructed and analysed the mitochondrial genomes of 60 individuals including 3 *Exaireta spinigera* (used as an outgroup). The 60 reconstructed mitochondrial genomes have the typical structure of insects mtDNA : 13 protein-coding genes, 22 tRNA, 2 rRNA-coding genes as well as non-coding regions and control regions (an example is given as a supplementary (SUPP_Figure 004 (SUPP_MATERIAL)). The genetic differences are homogeneously distributed on the whole mtDNA and do not seem to be localized on precise genes (Figure 5A). A great genetic diversity at the mitochondrial genome level is observed. Two very homogeneous groups A and C are well represented, and sequences of the C haplotype are very divergent all along the genomes from the others. The two sequences with the most differences are 55-F-Angouleme and 21-Ghana with a similarity percentage of 96.373% (all results are in supplementary (SUPP_Figure 005 (SUPP_MATERIAL)).

**Fig 5.**
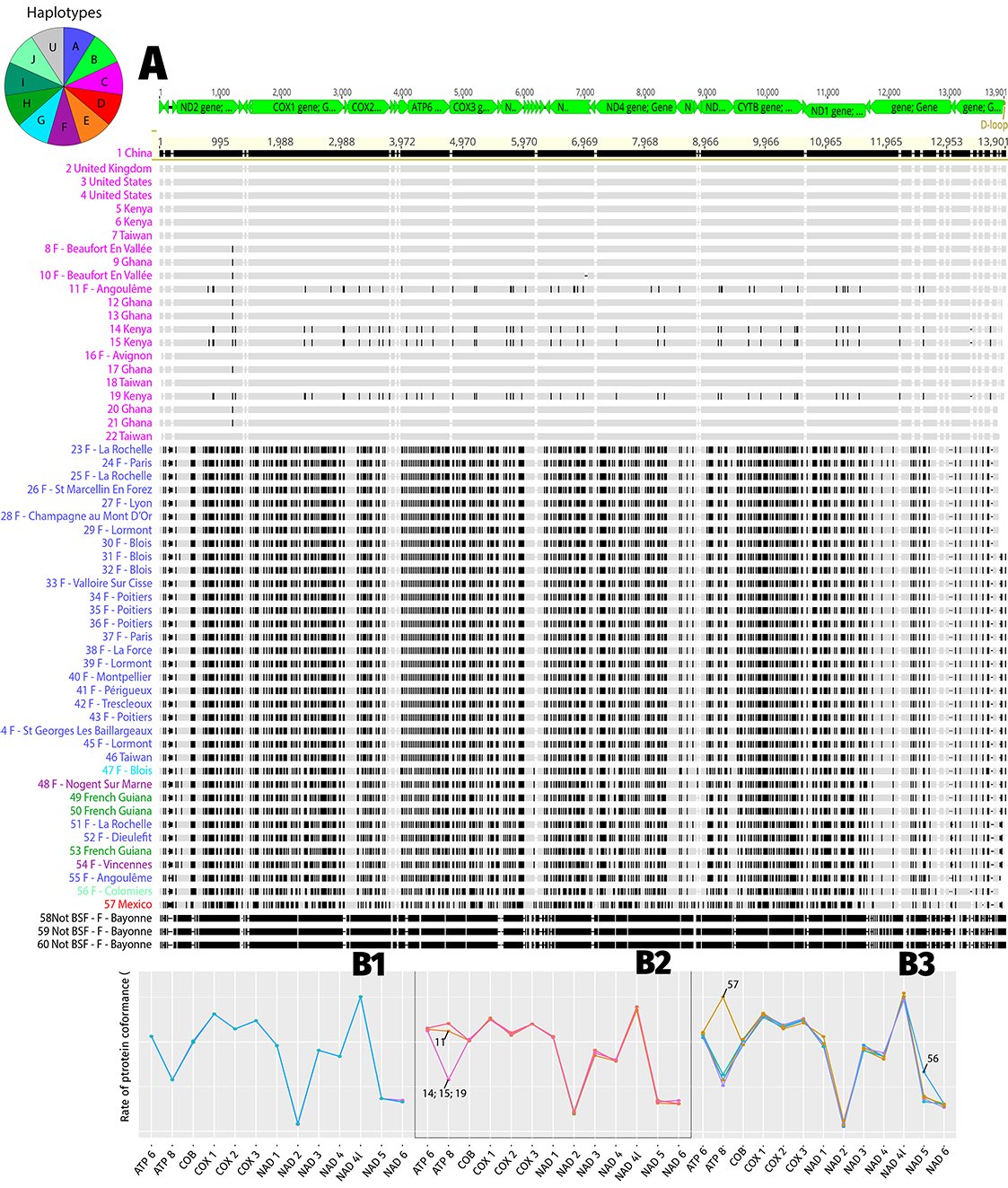
**A** Multiple alignment of mitochondrial DNA sequences from *Hermetia illucens*. Alignment performed with the MAFFT aligner. The colors of the names come from their haplotypes determined with the CO1 sequences. Each black mark represents a difference with the reference (1 – China). Above the alignment there is a representation of the mitochondrial genes. **B** Sequence conformity of mitochondrial genes of *Hermetia illucens* to *Exaireta spinigera*. The sequences are grouped according to haplotypes. The A haplotype (B1), the C haplotype (B2) and the other haplotypes (B3). The sequences inside each graph are those represented in the alignment (5A). Sequences from 23 to 46 plus sequences 51, 52, and 55 for haplotype A (5-B1); from 1 to 22 for haplotype C (5-B2); and from 47 to 57 (except sequences 51, 52 and 55) for the other haplotypes (5-B3).1

We also wanted to know if the mutations are equally (or not) distributed between the different genes. Therefore, we compared the level of similarity of the 13 mitochondrial genes of the 57 BSFs (fig 5B) to those of *Exaireta spinigera*. The results are presented as percent sequence similarity.

We grouped the individuals into 3 different categories:

– Haplotype A (Figure 5 B1), here all genes show the same differences with those of *Exaireta spinigera* only the NAD6 gene shows slight differences.
– Haplotype C (Figure 5 B2), we can see that most of the mitochondrial genes are identical except for the ATP8 gene which is clearly different for four individuals (11 Angouleme; 14 Kenya; 19 Kenya and 15 Kenya - Table available in supplementary 4 material). Slight differences are also present in the NAD3 and NAD6 genes.
– Haplotype D-F-G-H-I (Figure 5 B3), the profiles are all slightly different, with a stronger disparity for the ATP8, NAD1 and NAD5 genes, especially the Mexican one (57). The profiles show the same pattern except for the ATP8 gene of the individual from Mexico (57) which is more conserved.

### Complete mitochondrial genome phylogeny

The complete mitochondrial genome phylogeny demonstrates the very high level of genetic divergence among BSF populations that were found with the CO1 gene analysis (Fig 6A). The increased sensitivity of the analysis, due to the size of the sequences (13.96 kb, 13 genes) robustly resolves the relationships between different haplotypes identified previously (nodes >95%). In addition, it allows us to distinguish subgroups within the previously determined haplotypes. Two major divisions seem to emerge within the tree: group C (which includes all commercial strains) and all others. Within the C group two subgroups emerge (11 – F – Angouleme; 14 Kenya; 19 Kenya; 15 Kenya and the others).

**FIG 6.**
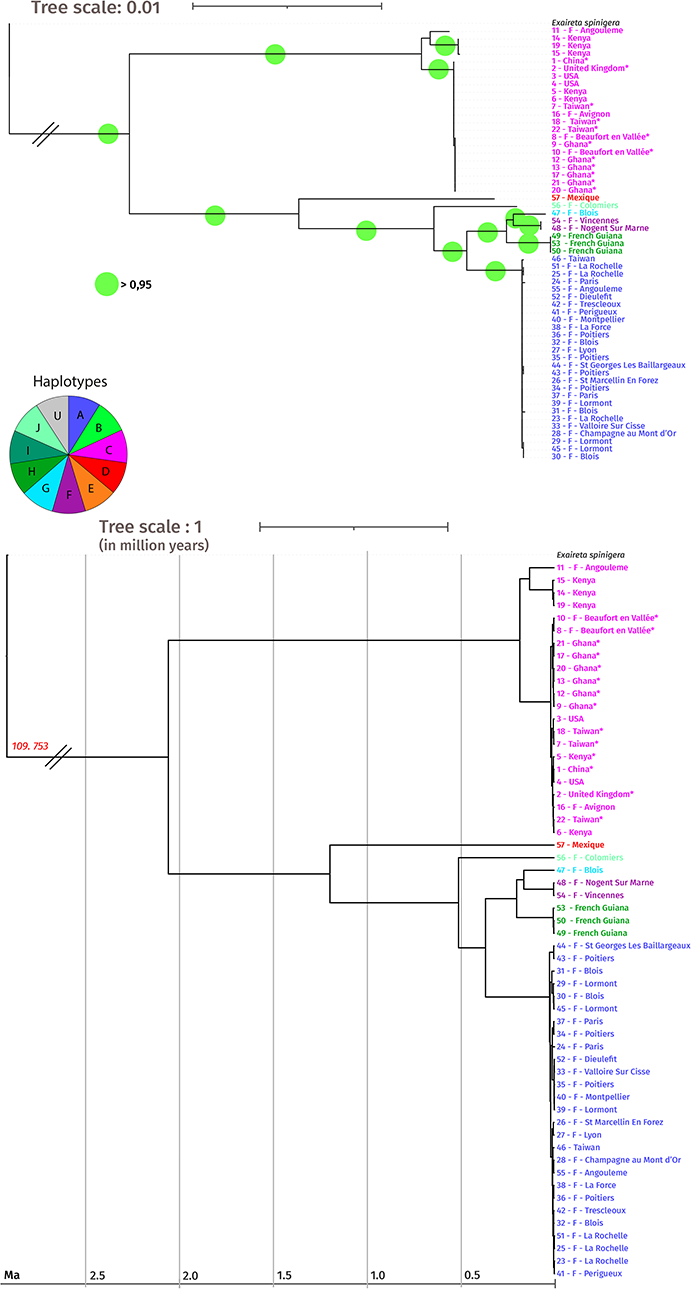
**A** ML phylogenetic tree, of 57 mitochondrial DNA sequences of *Hermetia illucens*, deprived of D-Loop regions. MAFFT was used to align the sequences, 500 Bootstrap replicas were performed. The outgroup used is *Exaireta spinigera* (Stratiomyidae). The colors of the names correspond to the haplotypes determined with the CO1 gene sequences (Figure 1). Bootstrap values are indicated by a green circle when they are > 0.95. Commercial individuals are marked with an asterisk. **B** Bayesian time tree obtained with BEAST of 60 mitochondrial genomes of *Hermetia illucens* rooted with *Exaireta spinigera*. The scale is in millions of years. A break in scale is made at the root level. The colors of the names correspond to the haplotypes determined with the CO1 sequences (figure 1). Commercial individuals are marked with an asterisk.

The other large group contains the sequences of the A D I G and H haplotypes. The I G and H haplotypes are related and close to each other. The very cosmopolitan A haplotype is well conserved with a limited level of mitochondrial sequence divergences.

Moreover, the I G and H haplotypes, although phylogenetically close, are found in very disparate areas (the G haplotype is mostly in Africa, the I in Europe, and the H in Latin America) supporting a multiple worldwide introduction.

Strikingly the high level of the conservation of the ATP8 sequence described above in the Mexican sample (haplotype D) does not explain the basal position of the sequence in the phylogeny; indeed, we tested the phylogeny by removing the ATP8 gene and no change was found (SUPP_Figure 006 (SUPP_MATERIAL)).

The ATP8 gene alone is not sufficient to explain the appearance of the observed subgroups. However, unlike the CO1 gene which varies only slightly within this haplotype, the NAD2, NAD3, NAD5 and NAD6 genes are more variable.

The average percentage of identity between haplotypes A and C is about 96.39%. Between haplotype C and D 96,54%. Between haplotype C and H 96.40% and 96.43% between C and I. Between A and D 97.82% (all results are in SUPP_Figure 005 (SUPP_MATERIAL)).

### Divergence time of the different groups

The time tree (fig 6B) gives us information about the divergence time of the different mitochondrial genomes. We obtained identical Bayesian and ML trees. We noted a relatively long divergence time between haplotype C and the others (2.2 million years). The others have diverged more recently (about 1.2 million years). Within the C haplotype, a divergence has been about 300.000 years while in the other groups the divergence times are much lower, about 20.000 years for the A haplotype.

The time of divergence with *Exaireta spinigera* has been estimated at 110 million years, which is confirmed by the literature that gives us 112 million years with the TIMETREE tool (Hedges et al, 2015).

## Discussion

*Hermetia illucens* is now considered as a species of major interest for the food industry (Tomberlin and Huis 2020). Thus, it became urgent to study the genetic diversity within the different populations at different scales. The analysis of the different haplotypes in the same geographical area is a powerful tool to study the recent history of BSF. An in-depth study of the genetic variations between the different haplotypes may allow us to better resolve the scenario of the arrival of BSF in a territory.

In this study, we uncovered a high genetic variability of mitogenomes within the species *Hermetia illucens*. We found that the haplotype C is the most divergent. The dating of the separation events with the other haplotypes goes back more than 2 million years. We found a well resolved separation within haplotype C (>95%); the other haplotypes are also well resolved (>95%).

Our study confirms the genetic diversity that was found in previous papers (Kaya et al, 2021 ; Park et al, 2017 ; Ståhls et al 2020). We found a comparable number of genetic clusters although we add many more sequences of diverse origin. For example, Ståhls et al, found 56 distinct haplotypes with 418 CO1 sequences and we found a comparable number of haplotypes based on the analysis of 677 CO1 sequences. These results suggest that we have now captured a major part of the extant BSF worldwide diversity. However in South America we found 30 haplotypes among the 55 sequences sampled, that open-up the possibility of the existence of a still unknown genetic diversity in this part of the world.

With the greatest genetic diversity being found in South America, the hypothesis of the geographical origin of BSF in Latin America is further confirmed. Apart from this continent of origin, there seem to be a huge number of neo-introductions on all continents with divergent mitochondrial lineages. The analysis of complete mitogenomes supports the hypothesis of several distinct introductions in Europe as well as on other continents. Our results contradict the scenario of Kaya et al based on microsatellites for a single BSF introduction in Asia and Australia. Indeed, our data support the presence of highly divergent major haplotypes A, C and E in Australia and major haplotypes A, B, C in Asia in addition to several orphan haplotypes. As the divergence times between these major haplotypes is estimated around two million years, this pattern is incompatible with a single introduction by man in the recent period but supports multiple and independent introductions. In the same way, Kaya et al. proposes at least three distinct introductions in Europe. Our data considerably increases this number: at least seven different haplotypes are found (A, I, J, G, F, C and E) supporting at least seven distinct introductions. Similarly, at least four introductions in Africa arose. By increasing the sample size and using more resolutive markers, we obtained a more precise description of BSF genetic diversity and a better understanding of its natural history. However, due to the frequent, multiple, and probably ongoing introductions worldwide of the BSF, the understanding of the exact introduction history might be very challenging.

Even at a national scale, the BSF genetic diversity is huge. By taking the example of France, we were able to show that the diversity observed at the global scale is also found at the national scale (five of the 10 major haplotypes found worldwide). Moreover, at the local level, we have haplotype mixtures, confirming the observations made on microsatellite data (Kaya et al, 2021). For example, in the region of Paris we have a mix of three haplotypes (A, F and I), and in the region of Montpellier, a mix of four haplotypes (A, B, C and E) – Figure 4.

France contains on its territory a great diversity of haplotypes, whose origin can be explained either by the historical introduction of the year 1950 (Barbier 1952), in addition to more recent neo-introductions. The haplotype A, which is by far the most common haplotype in wild BSF found in France (71.1%) and in Europe, may correspond to the initial introduction. However, it is difficult to understand the exact scenario of the introduction of BSF in France: for example, the divergence times within French flies belonging to haplotype A (20.000 years) tend to prove the recent but multiple introductions in France of flies belonging to this group rather than a single introduction by men followed by a diversification on the territory. Our data supports a complex scenario involving multiple and repetitive introduction of the BSF from closely related (same phylogenetic groups) and distantly related (divergent phylogenetic groups) individuals in France. A possible scenario involves an initial introduction of the haplotype A in the 1950s (Barbier 1952) which spread gradually on the territory. More recent introductions arose explaining the occurrence of the haplotypes J, G and I. Finally, introductions related to the industrial activity of the BSF of the haplotype C arose.

Jefferey Tomberlin, a pioneer in the field, also indicates in a recent article on the structure and demography of the BSF based on microsatellite data (Kaya et al, 2021), that the captive populations used for breeding in Europe and North America derive from a common, genetically related strain. Our analysis based on CO1 and complete mitochondrial genome sequences confirm this result: the haplotype that we have named C (for commercial) contains all the sequences of industrial origin with very few genetic diversity. Interestingly, this commercial group does not have any South American relatives but Kayla et al. reports wild north American populations that are related (but not similar) to the haplotype C. This result suggests a possible North American origin for the BSF commercial stock. Our data supports a possible alternative scenario because both network haplotypes and CO1 phylogeny indicate that the C haplotype is closely related to the B haplotypes, as well as 8 other haplotypes distributed in Asia and Europe. However, the exact origin of the commercial stocks might be difficult to decipher due to the naturalisation of the C haplotypes in many locations.

Indeed, certain individuals collected in the wild and belonging to this group were collected in the vicinity (around 5 km) of industries breeding BSF. For example, if we focus on the commercial haplotype analysis in France (the C haplotype), it is possible that the wild BSF belonging to the haplotype C have escaped from industries in the regions of Poitiers, Bordeaux, Avignon, and Montpellier. This last point raises the question of the biosecurity of BSF farms around the world. Indeed, if the industries carrying out breeding of BSF do not opt for an optimal safety, the escape of BSF of industrial origin can lead either to an establishment of the BSF in territories where it was not yet present, or to a replacement-hybridization of the local populations by the industrial strain. This would lead inexorably to a loss of the local genetic diversity thus potentially erasing some local adaptation to specific environmental conditions.

However, it is interesting to note that in Ståhls et al 2020, in contrast to Kaya et al 2021 and our study, some genetic diversity in the breeding strains in Europe and North America have been evidenced (10 haplotypes). This may be due to a broader sampling effort of commercial strains. Indeed, STAHLS had the opportunity to sample many small farms leading to 292 BSF sequences from rearing cultures. This allowed them to access a broader genetic diversity not found in our study and in Kaya et al. In addition, it is also possible that smaller farms did not obtain BSF from the dealers who constituted most of the larger industries, but directly from *in Natura* acquisitions. Nevertheless, our results confirm that the industrial actors work with the same and unique strain.

We also evidence through this study that the mitochondrial genome is perfectly suited to differentiate haplotypes and to resolve the phylogenetic relationships between them. It allows us to date the natural history of the BSF, bringing a sum of details that were impossible to distinguish based only on the CO1. For example, we report puzzling variations in the level of conservation of the ATP8 genes for some samples (haplotypes C and D). Interestingly, these variations have already been found in other insect species (for example in coleoptera see Zhang et al 2019, and in grasshoppers see Li et al 2018). The ATP8 gene is a small gene (160 bp on average in insects) that encodes a subunit of the complex V of the ATP synthase. In *Hermetia illucens*, the initiator codon of the ATP8 gene is ATT, is 167 bp long and it overlaps with the ATP6 gene. It has been assumed in other species that the modification of the sequence of the ATP8 gene can affect the structure of mitochondrial complex V and thus its function. Furthermore, it has been shown that changes within ATP synthase can be explained by adaptation to different ecological environments (Peng et al, 2019 ; Gu et al, 2016 ; Zhang et al, 2017). This indicates that some local adaptation driven by the ATP8 genes may exist in BSF, as it has already been shown in other insects. This can lead to an adaptation to survive in high altitude environments, i.e., at a lower level of oxygen and low temperature (Luo et al, 2013). Even if these variations did not change the phylogeny of the BSF it can be a clue to some local adaptation signal.

Complete mitochondrial genome data allows us to make a robust and well resolved time tree of the species. The divergence times between the different haplotypes are surprisingly long (more than two million years for the most divergent lineages) and the differences observed between the sequences tend to prove that there is still unsuspected and hidden genomic diversity for the species. If we compare it to Drosophila, two million years correspond to many speciation events leading to different reproductively isolated species (Russo et al, 2013). By contrast, Stahls et al indicated that BSF belonging to phylogenetically divergent haplotypes are still able to cross and generate offspring, indicating that there is no apparent reproductive isolation in laboratory conditions. Interestingly, our data evidence the cohabitation of different haplotypes at the same locality that could lead to crosses between haplotypes. Further investigations are thus needed to understand the possible genetic barriers between highly divergent BSF groups but also the possible patterns of introgression in natural BSF populations. This phenomenon might considerably confuse the elucidation of the population structure and the natural history of these species.

Finally, it is interesting to recall that, if indeed, most of the industrial actors use the same haplotype for the industry, the other haplotypes present in France are very divergent from this one. Strikingly, we found in one compost bin in the region of Paris flies belonging to two different haplotypes (A and F). Thus, crossbreeding between these two haplotypes would bring a strong genetic mix that could be beneficial for breeding. Associated with this significant genetic diversity, BSF may also have some significant phenotypic diversity that might be useful for specific industrial usage and selection processes to increase performances or to ameliorate peculiar traits or behaviours (Zhou et al, 2013).

## Conclusion

During this study, through the analysis of 60 mitochondrial genomes, we were able to propose a robust and well resolved time tree of the BSF that confirms the huge and hidden genetic diversity within the species *Hermetia illucens*. Our result elucidates the phylogenetic relationship and time divergence between the different BSF lineages and provides a plausible scenario for BSF expansion in the world via multiple and independent introduction at the global and local scale. With the use of 677 COI gene sequences coming from a broad geographical sample, we were able to highlight the genetic diversity at the level of the different continents evidencing recurrent and multiple introductions outside the geographical area of origin in Latin America. However, the exact scenario of introductions, even at a national scale in France, are still difficult to understand due to probable multiple and independent introductions in addition to possible interbreeding between different strains that coexist in the same geographical area. We have also shown that a large part of the BSF industrial companies work with flies that derive from a unique strain. We would also like to remind that the apparent biosecurity deficiency within some BSF industrial companies probably led to escapement and possible local introduction/replacement of natural populations of BSF. Finally, our data indicated that it becomes extremely interesting to take a closer look at the nuclear genomes’ evolution of the BSF to better understand the natural history of the BSF as well as specific adaptations of the different lineages that might be useful for industrial needs and to initiate the selection process for peculiar traits.

## Availability of data and materials

The dataset generated during the study has been deposited in the European Nucleotide Archive database under Accession Numbers PRJEB48031 Bioprojet. Alignment and raw data can be downloaded following this link http://gofile.me/2ppPR/QWjM33T7y the supplementaries can be downloaded following this link : http://gofile.me/2ppPR/rwaGSXDQU

## Contributions

JG carried out the sampling, performed data curation, and wrote the original draft. NP performed data curation and reviewed the manuscript. GB helped with the data curation, and reviewed the manuscript. JF conceptualised and supervised the study, performed data curation, reviewed, and finalized the manuscript.

## Acknowledgements

We would like to thank Floran Laville and Frederic Marion-Poll for his continuous support and help during this project. We warmly thank the members of the Réseau Compost Citoyen for their precious help during the sampling in France. We also thank Matan Shelomi (National Taiwan University), Meng-Kun Wu, Shu-Min Chen, Jing-Jiun Huang (Yi Mi Community College, Taiwan) for flies from Taiwan. We thank the international Centre of Insect Physiology and Ecology (ICIPE) from Nairobi City, Kenya. We thank Mathieu Chouteau from CNRS LEESIA, in French Guiana. We finally thank Benoit Gilles for his help at the beginning of this project.

